# METTL3 inhibitors for epitranscriptomic modulation of cellular processes

**DOI:** 10.1101/2020.09.25.311803

**Authors:** Elena V. Moroz-Omori, Danzhi Huang, Rajiv Kumar Bedi, Sherry J. Cheriyamkunnel, Elena Bochenkova, Aymeric Dolbois, Maciej D. Rzeczkowski, Lars Wiedmer, Amedeo Caflisch

**Author notes:** To whom correspondence should be addressed.; Correspondence may also be addressed to Elena V. Moroz-Omori. Elena V. Moroz-Omori and Danzhi Huang contributed equally.

## Abstract

The methylase METTL3 is the writer enzyme of the N^6^-methyladenosine (m^6^A) modification of RNA. Using a structure-based drug discovery approach, we identified a METTL3 inhibitor (**UZH1a**) with potency in a biochemical assay of 280 nM, while its enantiomer **UZH1b** is 100 times less active. The crystal structure of the complex of METTL3 with **UZH1a** illustrates the interactions that make it selective against protein methyltransferases. We observed a dose-dependent reduction in m^6^A methylation level of mRNA in several cell lines treated with **UZH1a** already after 16 h of exposure, as determined by triple-quadrupole LC mass spectrometry, while its enantiomer **UZH1b** was essentially inactive at concentrations up to 100 μM. Interestingly, the kinetics of m^6^A level reduction in mRNAs followed a first-order reaction model, with a half-decay time τ of 1.8 h and a maximum m^6^A inhibition level of 70%, which is in line with the previously observed shorter half-life of m^6^A-modified mRNAs. Notably, treatment with the compounds did not alter cellular METTL3 levels, ruling out indirect effects on m^6^A levels. The effect of the m^6^A level depletion by **UZH1a** directly translated into growth inhibition of MOLM-13 leukemia cells, under short-term and long-term culture. Incubation of the MOLM-13 cells with **UZH1a**, but not with **UZH1b**, resulted in increased cell apoptosis and cell cycle arrest already after 16 h of incubation. Interestingly, other cell lines sensitive to METTL3 level (U2Os, HEK293T) did not reveal statistically significant differences between **UZH1a** and **UZH1b** in a cell viability assay, confirming that the degree of reliance on m^6^A signalling for survival can vary between cancers/cell types.

## Introduction

The finely organized network of gene expression comprising RNA transcription, splicing, transport, translation, and degradation is often perturbed in cancer.^1,2^ In addition to previously known regulators of gene expression, such as epigenetic modifications or miRNAs, the recently discovered layer of regulation based on co- and post-transcriptional RNA modifications gave rise to a new field named epitranscriptomics.^2,3^ While over 160 different RNA modifications have been discovered to date, one of the most abundant modifications, N^6^-methyladenosine or m^6^A (comprising 0.1 – 0.4% of all adenosine in mRNA), is involved in most of the aspects of RNA regulation, *i.e.*, alternative polyadenylation, splicing (controls about 3% of alternatively spliced exons^4^), nuclear export, stability, and translation initiation (**Fig. 1**).^2,3^ It is also found in other RNA species including lncRNAs,^5^ rRNAs,^6^ and snRNAs.^7^ Dysregulated m^6^A deposition is directly involved in the development of acute myeloid leukemia (AML) and lymphomas, difficult-to-treat blood cancers, ^1,8–12^ as well being associated with other types of cancer (*e.g.*, bladder, lung, ovarian, colorectal, bone, liver, gastric).^13–21^

**Figure 1.**
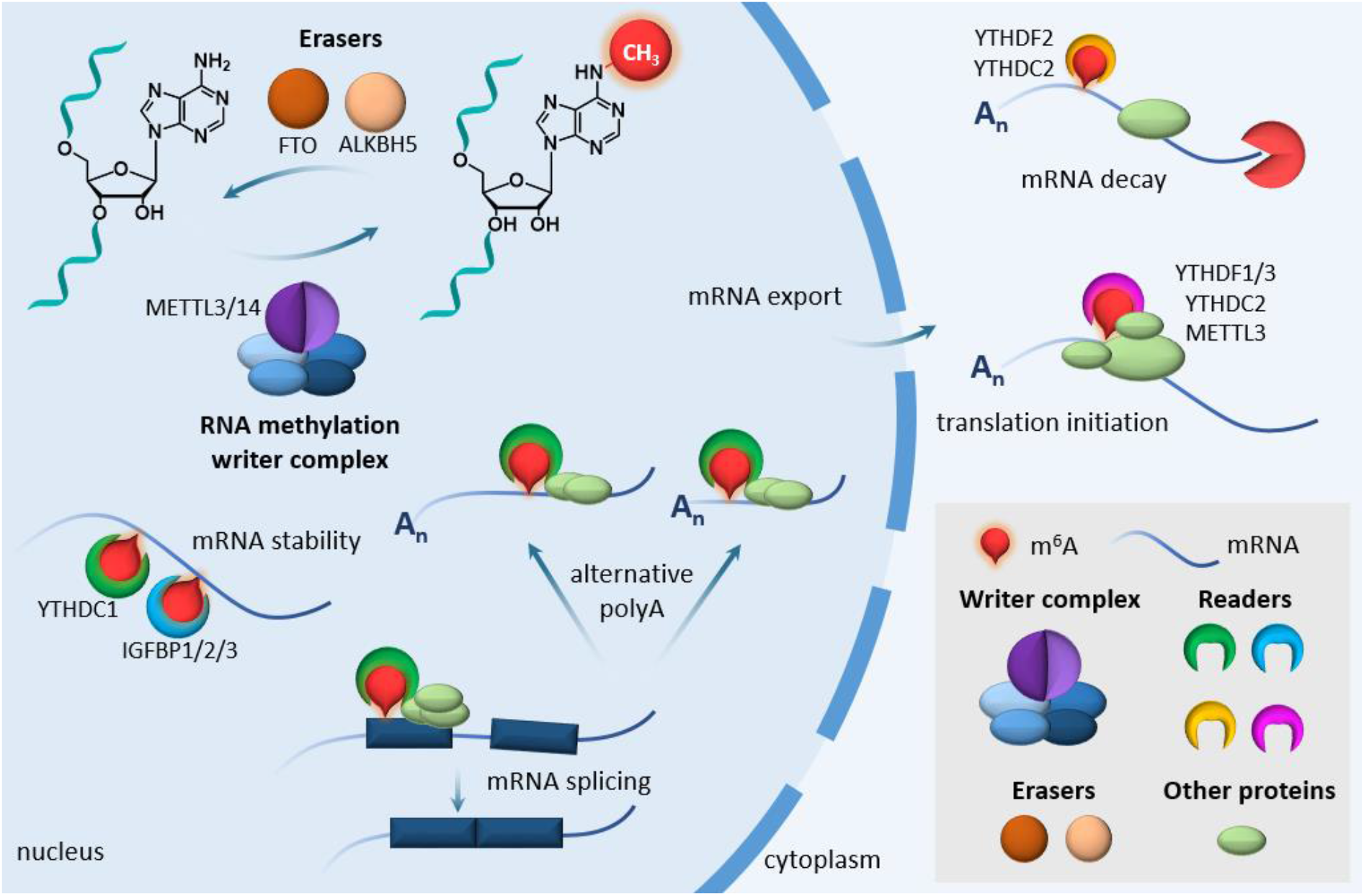
The m^6^A methylosome. Schematic representation of m^6^A regulatory proteins and diversity of their functions.

In mRNAs and lncRNAs most of the m^6^A modifications are installed in [G/A/U][G>A]m^6^AC[U>A>C] consensus sites (GGm^6^ACU is the most prevalent)^22^ by the so-called m^6^A-METTL complex (MAC), consisting of methyltransferase-like protein 3 (METTL3) and METTL14.^3^ MAC is assisted by a regulatory complex (named MACOM, <u>m</u>^6^<u>A</u>-METTL-associated <u>com</u>plex) composed of WTAP, RBM15/B, VIRMA, ZC3H13 and HAKAI.^23^ The crystal structure of the METTL3/14 complex was resolved in 2016,^24–26^ revealing the function of each component. While METTL14 plays a scaffolding role in substrate RNA recognition, forming an RNA-binding groove at the interface of the two subunits, METTL3 carries out the catalytic transfer of the methyl group from a cofactor *S*-(5′-adenosyl)-ʟ-methionine (SAM) onto the N^6^ atom of the adenine.^24–26^

METTL3/14 plays a critical role in disorders associated with abnormal self-renewal, aberrant proliferation, and blocking differentiation of hematopoietic cells.^1,8–12^ AML and B-cell lymphoma are among the cancers with the highest expression of METTL3 based on The Cancer Genome Atlas database.^9^ Both METTL3 and METTL14 were found to be upregulated in all subtypes of AML compared to normal hematopoietic cells, despite the heterogeneity of this blood cell cancer in terms of chromosomal rearrangement and gene mutations.^8^ More importantly, a genome-wide CRISPR-based screen to identify genes essential for cell survival using 14 distinct AML cell lines scored all members of the writer complex, including METTL3 and METTL14, among the top 10% of the essential genes, similar results were obtained in mouse leukemia.^9,27^ Strikingly, the apoptotic response is specific for leukemic cells and it has not been observed in normal hematopoietic progenitors.^8–10^ These phenotypes are at least in part due to METTL3 regulation of the translation of MYC, BCL2, and PTEN, which are central regulators of cell survival and differentiation. Inhibition of PI3K/AKT signaling partially reverses induction of differentiation in METTL3 knockdown cells.^10^ In addition, METTL3, independently of METTL14, binds to chromatin and is recruited to transcriptional start sites of a set of target genes highly enriched for CAATT-box binding protein CEBPZ binding motifs (*e.g.*, SP1 and SP2), thereby controlling their translational efficiency by reducing ribosome stalling due to the presence of m^6^A.^9^

To date, there are no cell-permeable inhibitors of METTL3/14 with a disclosed chemical structure except for the universal nucleoside analogue sinefungin, which inhibits most methyltransferases and is not selective for METTL3.^28,29^ Therefore, all published studies of the role of m^6^A in cancer have relied on the knockdown/overexpression of the writers, erasers, and readers. In contrast to this approach, using small molecule inhibitors preserves the function of the target enzyme to act as a scaffold for protein–protein interactions that would otherwise be disrupted by RNAi, thus enabling discrimination between the enzymatic and structural roles. Indeed, it has been demonstrated that both a catalytically active and a non-functional METTL3 lead to a reduction in p-AKT levels highlighting the m^6^A-independent function of METTL3.^10^ In addition, small molecule inhibitors enable titration experiments across a range of concentrations to be performed, ranging from mild perturbation to near complete inhibition of the METTL3 enzymatic activity, which can reveal a spectrum of phenotypes. Generally, inhibitors can inactivate their targets rapidly, allowing precise temporal control over METTL3 function. This is particularly important for studying downstream effects of m^6^A depletion on gene and protein expression and will help to elucidate the m^6^A regulatory network in health and disease.

Here we report a nanomolar inhibitor of METTL3 (**UZH1a**) which is selective and cell-permeable, while its enantiomer **UZH1b** is essentially inactive. The crystal structure of the complex shows several favourable interactions of the **UZH1a** chemical probe with METTL3, some of which are not observed in the METTL3/SAM complex and thus provide selectivity against other SAM-dependent methyltransferases. Furthermore, the characterization of **UZH1a** in biochemical and cellular assays provides evidence that **UZH1a** has a good potential for the development of a lead compound against AML.

## Results and Discussion

Our initial efforts to develop a potent and selective METTL3 inhibitor were focused on screening an adenine-based library, which yielded several hits with micromolar potency in a homogenous time-resolved fluorescence (HTRF) enzyme inhibition assay.^30^ Protein structure-based optimization coupled with compound potency evaluation in our HTRF assay^31^ resulted in the high-nanomolar inhibitor **UZH1a** (**Fig. 2a**). The potency of **UZH1a** in the HTRF assay was 280 nM, while its enantiomer **UZH1b** was 100-fold less active (**Fig. 2b**). We confirmed the binding of **UZH1a**, but not its enantiomer **UZH1b**, to METTL3 by X-ray crystallography (**Fig. 2c**). The **UZH1a** inhibitor fills the pocket of the adenosine moiety of SAM but not the pocket of the SAM methionine. Besides the favourable van der Waals contacts, there are several hydrogen bonds between polar groups of the **UZH1a** inhibitor and METTL3 (yellow dashed lines in **Fig. 2c**). There is also an intra-inhibitor hydrogen bond between the amide NH and phenolic oxygen of **UZH1a** (green dashed line in **Fig. 2c**). Notably, the binding of **UZH1a** results in a conformational rearrangement (6 Å displacement of the amino group) of the Lys513 side chain with respect to the Lys513 orientation in the complex of METTL3 with SAM or sinefungin (**Fig. 2d**, PDB code 6Y4G). In detail, the tertiary amino group of **UZH1a** replaces the primary amino of Lys513 in the salt bridge with Asp395. As a consequence, the amino of Lys513 replaces the amino group of SAM in the salt bridge with Glu532. These castling-like changes of ligand/METTL3 and intra-METTL3 ionic interactions are likely to provide selectivity against other SAM-dependent methyltransferases. Indeed, **UZH1a** demonstrated high selectivity towards other protein methyltransferases (as well as several promiscuous protein kinases), with remaining enzymatic activity of over 75% at 10 μM concentration of **UZH1a** (**Table 1**).

**Table 1.** Kinases and protein methyltransferases selectivity profile of **UZH1a** at 10 μM.

| Type | Protein | Remaining activity, % |  | Control IC<sub>50</sub> (µM) | Control compound |
| --- | --- | --- | --- | --- | --- |
|  |  | Replicate 1 | Replicate 2 |  |  |
| methyltransferase | DOT1L | 100 | 98 | 0.278 | SAH |
|  | G9a | 96 | 99 | 1.95 | SAH |
|  | MLL4 complex | 88 | 89 | 2.11 | SAH |
|  | PRDM9 | 101 | 101 | 3.98 | chaetocin |
|  | PRMT1 | 91 | 88 | 0.140 | SAH |
|  | SETD2 | 99 | 103 | 7.39 | SAH |
|  | SMYD3 | 88 | 92 | 18.7 | SAH |
| kinase | Abl | 80 | 74 | - | - |
|  | CDK2/cyclinA | 83 | 72 | - | - |
|  | cKit | 95 | 101 | - | - |
|  | DDR1 | 96 | 85 | - | - |
|  | Flt3(D835Y) | 80 | 68 | - | - |
|  | Fms | 92 | 102 | - | - |
|  | JAK1 | 101 | 102 | - | - |
|  | PDGFRα | 88 | 81 | - | - |
|  | Pim-1 | 100 | 95 | - | - |
|  | ROCK-I | 86 | 80 | - | - |

The remaining activity is the percentage of enzymatic activity in the presence of 10 μM **UZH1a** with respect to the buffer solution containing DMSO. The closer to 100% are these values, the weaker the inhibitory potency of **UZH1a**. SAH = S−(5′-adenosyl)-L-homocysteine.

**Figure 2.**
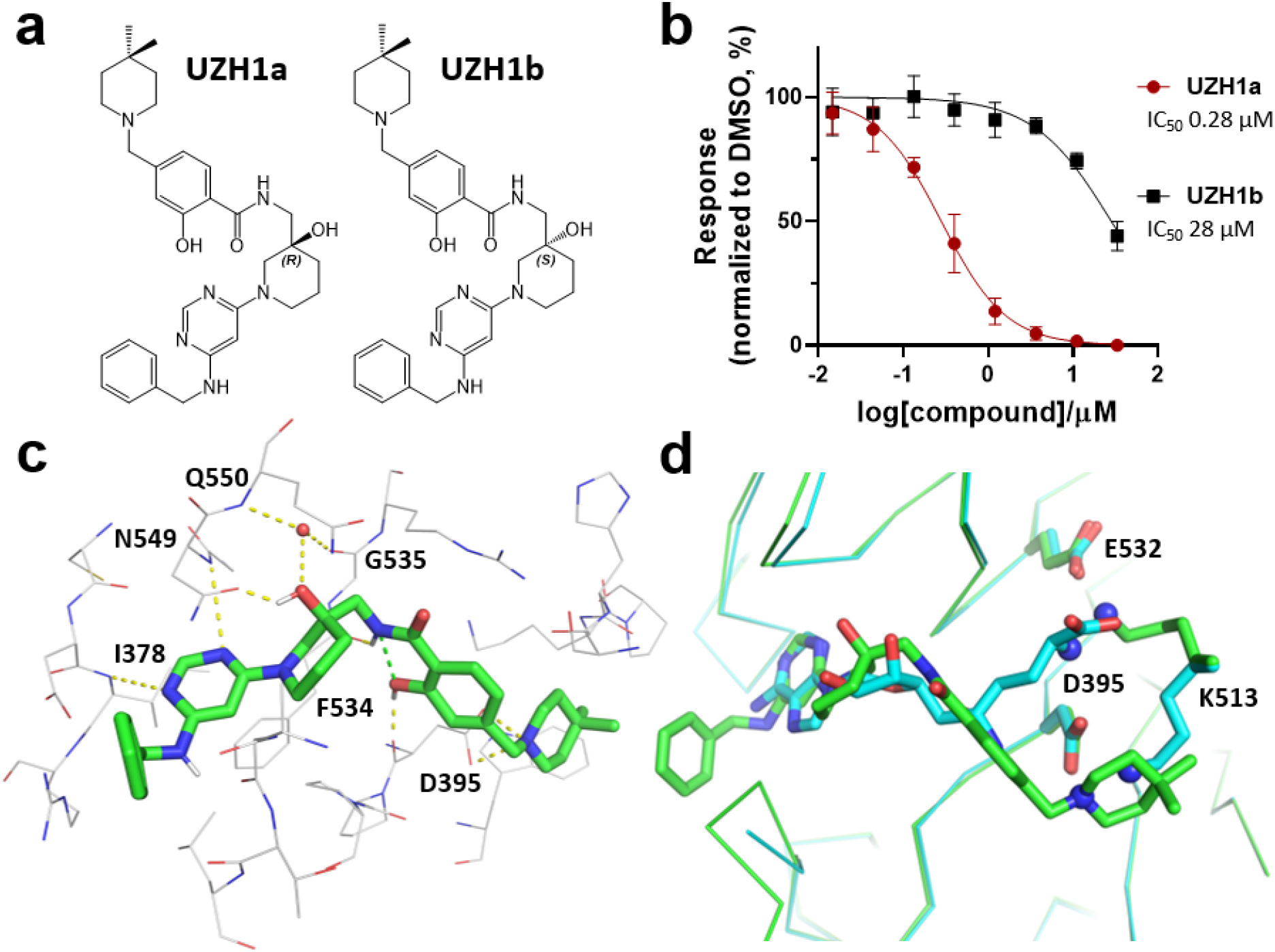
*In vitro* characterization of METTL3 inhibitors. (a) Chemical structures of **UZH1a** and **UZH1b**; absolute configuration was determined by X-ray crystallography. (b) Enzymatic activity assay based on HTRF, mean ± SD, N = 4. (c) Crystal structure of METTL3/14 in complex with **UZH1a** (carbon atoms in green, PDB code 7ACD). Hydrogen bonds (yellow and green dashed lines) and a water molecule in contact with the inhibitor (red sphere) are shown. (d) Overlay of crystal structures of METTL3/14 in complex with **UZH1a** (carbon atoms in green) and METTL3/14 in complex with sinefungin (carbon atoms in cyan, PDB code 6Y4G). The amino groups involved in the castling-like conformational change are highlighted (blue spheres on N atoms).

**UZH1a** possesses favourable physicochemical properties and is therefore compatible with cell-based experiments. Indeed, its molecular weight is relatively low (558 g/mol), and the octanol-water partition coefficient logD_7.4_ of 2.6 is optimal for cell uptake. **UZH1a** was highly permeable (P_app_ >1·10^−7^ cm/s) in a Caco-2 permeability assay used to evaluate intestinal uptake (**Table 2**). However, the efflux ratio of >2 suggests that **UZH1a** may be subject to active efflux. Importantly, the large difference in biochemical potency of **UZH1a** and **UZH1b** (> 100-fold, **Fig. 2b**) allows for discrimination of non-specific effects due to the chemical properties of the compounds and makes them highly suitable for mechanistic cellular studies.

**Table 2.** Permeability of reference compounds and **UZH1a** in Caco-2 cell assay.

| Compound | Direction | $P_{app}$ ($10^{-6}$ cm/s) | | $P_{app}(B-A)/P_{app}(A-B)$ |
| --- | --- | --- | --- | --- |
|  |  | Replicate 1 | Replicate 2 |  |
| Metoprolol | A-B | 33.70 | 31.37 | 1.05 |
|  | B-A | 34.71 | 33.32 |  |
| Atenolol | A-B | 0.48 | 0.42 | 2.04 |
|  | B-A | 0.89 | 0.95 |  |
| Erythromycin | A-B | 0.32 | 0.24 | 62.89 |
|  | B-A | 18.06 | 17.54 |  |
| **UZH1a** | A-B | 12.38 | 11.78 | 2.25 |
|  | B-A | 25.57 | 28.81 |  |

In line with the biochemical assay, we observed a dose-dependent reduction in m^6^A methylation level in mRNA from cells treated with **UZH1a** (IC_50_ of 4.6 μM), while **UZH1b** was less active at concentrations up to 100 μM in the human leukemia cell line MOLM-13, as determined by triple-quadrupole LC mass spectrometry (**Fig. 3a**). However, the effect of the m^6^A level depletion upon **UZH1a** treatment was attenuated by at least one order of magnitude in cells in comparison to the biochemical assay. This could be the consequence of cellular efflux (**Table 2**) or competition with high intracellular SAM/SAH levels. We confirmed that **UZH1a** was active not only in the leukemia cell line MOLM-13, but also in at least two other cell lines (*i.e.*, immortalized human embryonic kidney cell line HEK293T and osteosarcoma U2Os cells, **Fig. 3b**). We have verified that this effect was not caused by a reduction in METTL3 protein level (**Fig. 3d, e**). Interestingly, m^6^A level reduction in mRNAs followed a first-order kinetics, with a half-decay time τ of 1.8 h and a maximum m^6^A inhibition level of 70% (**Fig. 3c**), which is lower than the average mRNA half-life in cells (≈ 10 h)^32^ and is in line with the previously observed rapid degradation of m^6^A-modified mRNAs.^4,33,34^ Using **UZH1a** we could, for the first time, directly estimate the average half-life of m^6^A-modified RNA in the cells. This result illustrates an important advantage of chemical inhibition of METTL3 over genetic manipulation, as it allows precise temporal control over METTL3 function due to the rapid cellular penetration of small molecule probes. One has to bear in mind that this method does not allow the decay of m^6^A-modified mRNA to be distinguished from the potential demethylation of the transcripts *via* cellular ALKBH5. The latter, however, has been shown to affect only a small percentage of mRNAs after transcription.^4^

**Figure 3.**
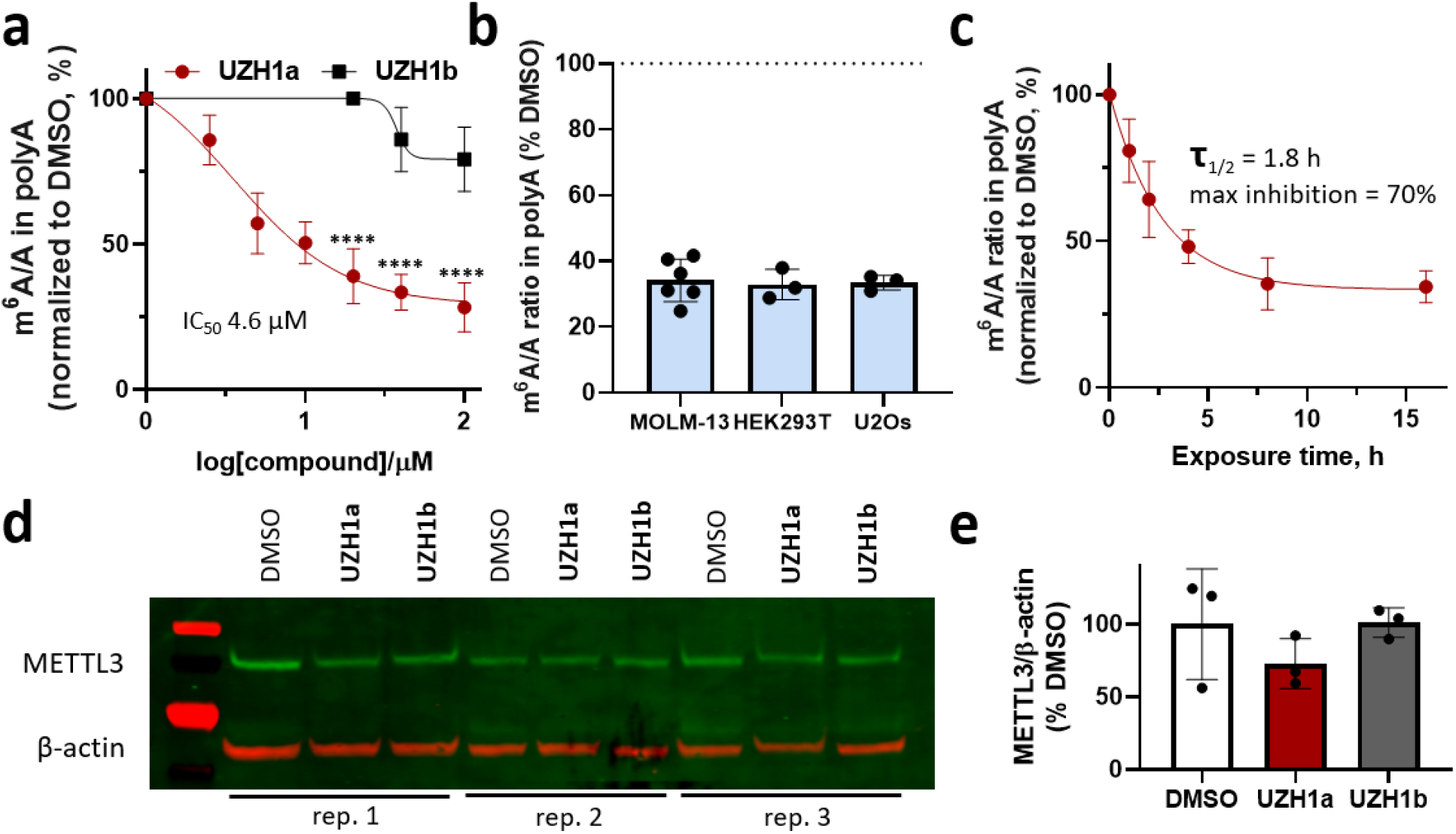
Cellular activity of the METTL3 inhibitor **UZH1a**. (a) Results of UPLC-MS/MS assay of m^6^A level in mRNA from MOLM-13 cells upon **UZH1a** and **UZH1b** treatment at a dose range of 2.5 to 100 μM for 16 h, mean ± SD, N = 3 – 7, \*\*\*\**p* < 0.0001 vs. **UZH1b**. (b) Results of UPLC-MS/MS assay of m^6^A level in mRNA from MOLM-13, HEK293T, and U2Os cells upon treatment with **UZH1a** at a dose of 40 μM for 16 h, mean ± SD, N = 3 – 6, normalized to DMSO-treated samples for each cell line. (c) Kinetics of m^6^A level reduction in mRNA from MOLM-13 cells upon **UZH1a** treatment at a dose of 40 μM for 1 to 16 h determined in UPLC-MS/MS assay, mean ± SD, N = 3, normalized to DMSO-treated samples for each time point. (d) Western blotting of METTL3 expression level in MOLM-13 cells after treatment with 40 μM of **UZH1a** and **UZH1b** for 16 h, N = 3. (e) Quantification of the METTL3 signal intensity relative to β-actin in western blot images of compound treated versus DMSO only treated samples, mean ± SD, N = 3.

Encouraged by the ability of **UZH1a** to decrease cellular m^6^A level, we further studied its efficacy in reducing proliferation of acute myeloid leukemia cells. We used as a model MOLM-13 leukemia cells as they have been extensively studied by the epitranscriptomics field due to their reliance on METTL3 for survival.^8–10^ In the short-term viability studies, the effect of the m^6^A level depletion by **UZH1a** directly translated in growth inhibition of MOLM-13 cells (GI^50^ = 11 μM), while its enantiomer **UZH1b** was 7-fold less toxic (**Fig. 4a**). The observed toxicity of the control compound **UZH1b** at high concentrations is likely caused by hydrophobic interactions with the cell membrane and/or off-target binding. A similar trend was observed in the long-term colony formation assay in MOLM-13 cells, where **UZH1a** significantly reduced the total colony area in comparison to **UZH1b** (**Fig. 4c, d**). This effect, however, was observed only at higher concentrations of **UZH1a**, presumably due to the presence of growth-stimulating factors in the colony culture medium. Reduced cell viability in presence of 20 μM of **UZH1a**, but not its enantiomer **UZH1b**, was in line with increased cell apoptosis already after 16 h of incubation, as evaluated by Annexin-V staining (**Fig. 5a, c**). The cell cycle analysis after 16-h compound treatment revealed that inhibition of METTL3 with **UZH1a** led to cell cycle arrest, as evidenced by the increased ratio of cells in G1 phase and decreased ratio of cells in S phase (**Fig. 5b, d**). Interestingly, other cell lines sensitive to METTL3 protein level (U2Os, HEK293T; **Fig. 4b**) did not reveal statistically significant differences between **UZH1a** and **UZH1b** in a cell viability assay (**Fig. 4a**), confirming that the degree of reliance on m^6^A signalling for survival can vary between cancers/cell types.

**Figure 4.**
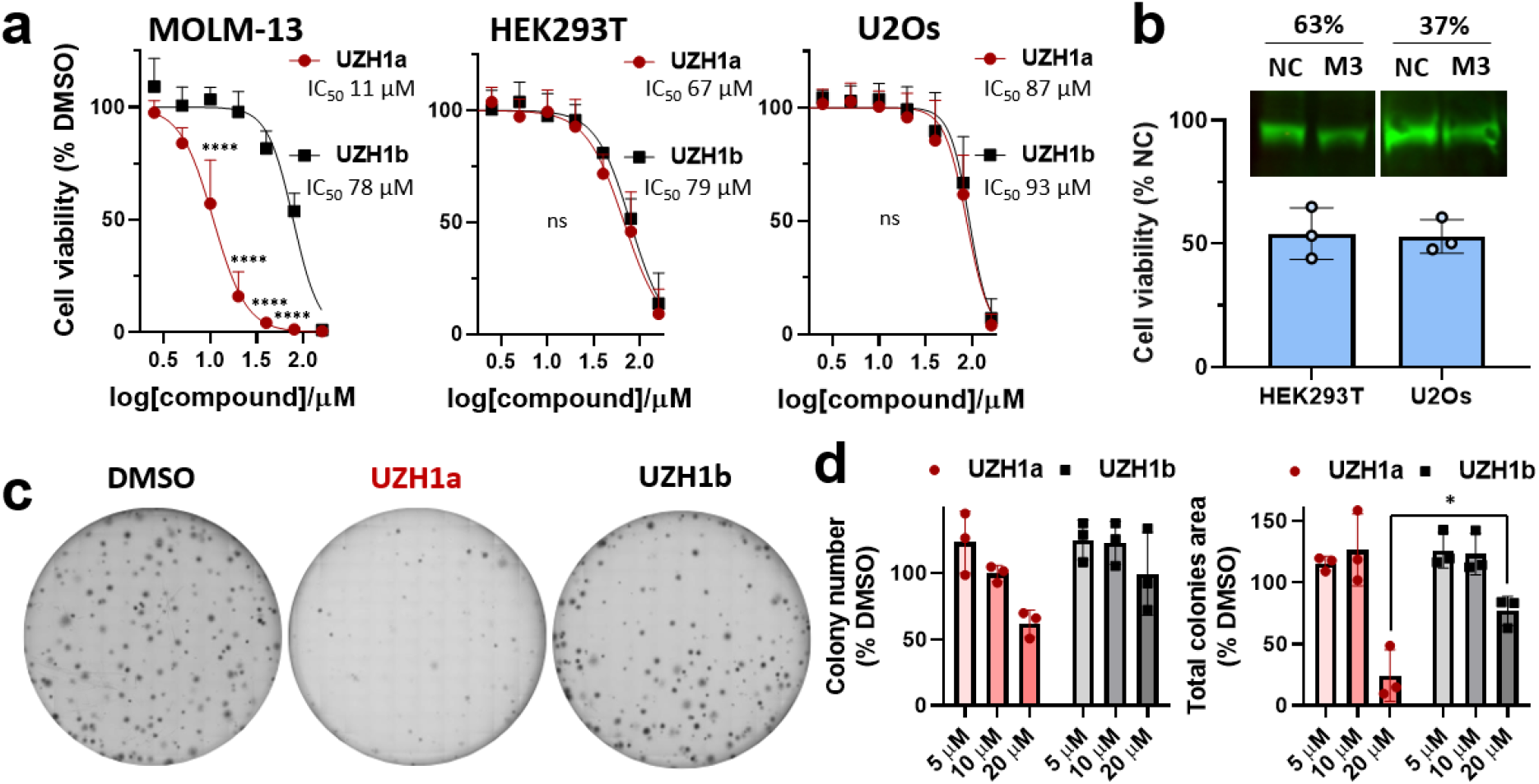
Effect of METTL3 inhibition on cell viability and clonogenicity. (a) Growth inhibition effect of **UZH1a** and **UZH1b** in MOLM-13, HEK293T, and U2Os cells after 72-h treatment, mean ± SD, N = 3 – 4, \*\*\*\**p* < 0.0001 in comparison to the same concentration of **UZH1b**, ns – not significant (*p* > 0.05). (b) Growth inhibition effect of METTL3 knockdown in HEK293T and U2Os cells after 72-h transfection with 50 nM of siRNA targeting METTL3 (M3) or a negative control siRNA (NC), mean ± SD, N = 3. Images above the cell viability plot represent corresponding results of western blotting analysis of METTL3 protein level reduction after the siRNA-mediated METTL3 knockdown. Numbers indicate residual METTL3 level in comparison to NC-treated samples. (c) Representative images of colony formation assay of MOLM-13 cells treated with 20 μM of **UZH1a** and **UZH1b** for 10 days in methylcellulose-containing medium. (d) Results of image analysis of colony formation assay of MOLM-13 cells treated with 5, 10, and 20 μM of **UZH1a** and **UZH1b** for 10 – 12 days, mean ± SD, N = 3, \**p* = 0.036.

**Figure 5.**
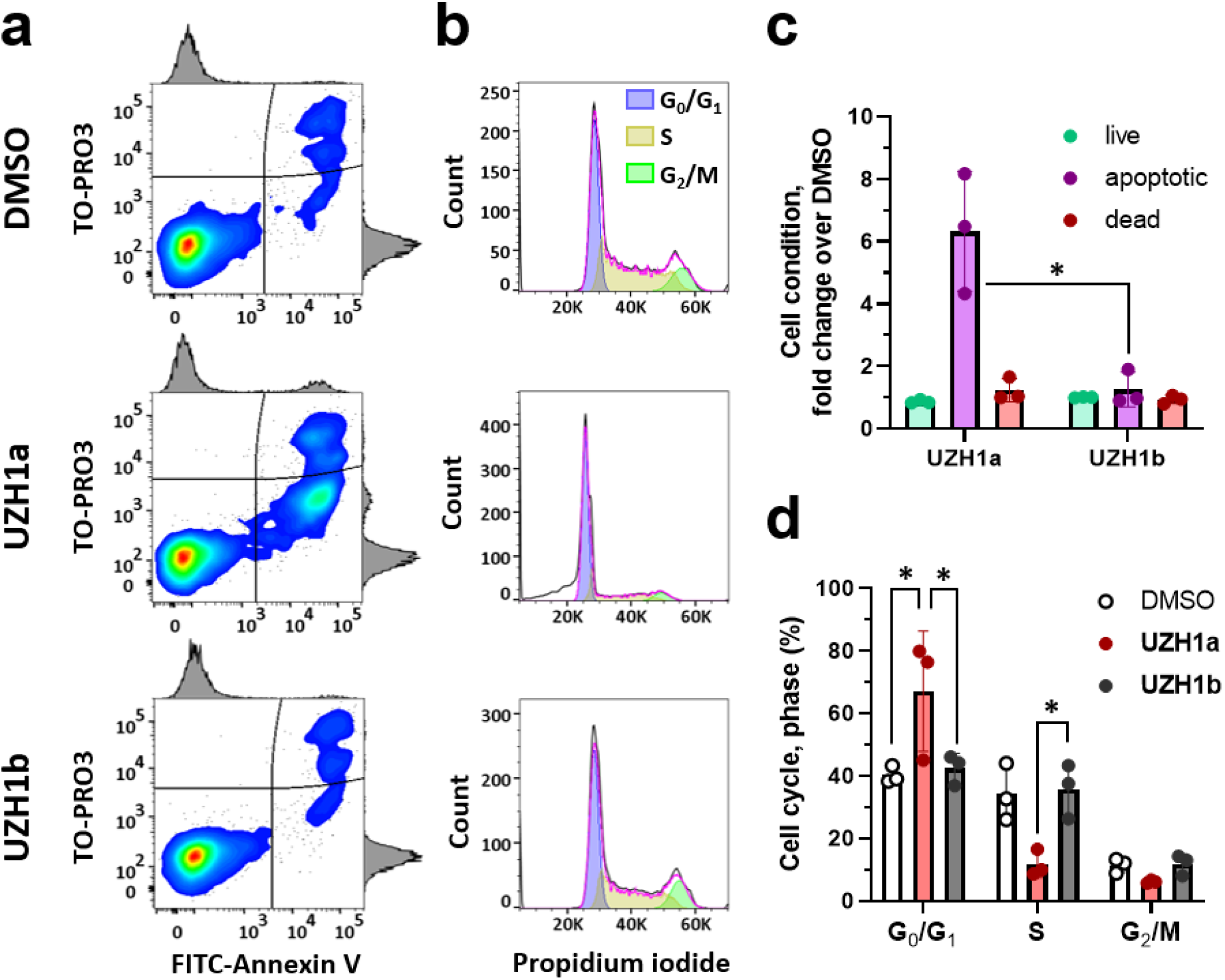
The effect of **UZH1a** and **UZH1b** on apoptosis and cell cycle in MOLM-13 cells. (a) Representative results of flow cytometry assay of FITC-Annexin V (bottom axis) and TO-PRO3 (left axis) stained cells after 16 h of exposure to 20 μM of compounds. (b) Representative results of cell cycle flow cytometry assay after 16 h of exposure to 20 μM of compounds. (c) Results of flow cytometry assay of apoptosis after 16 h of exposure to 20 μM of compounds, mean ± SD, N = 3, \**p* = 0.0257 (Kruskal-Wallis non-parametric test by ranks). (d) Results of flow cytometry assay of cell cycle after 16 h of exposure to 20 μM of compounds, mean ± SD, N = 3, \**p* < 0.05.

## Conclusions

Chemical probes that target proteins involved in the m^6^A modification of RNA are essential for understanding the complexity of this epitranscriptomic regulatory network. Furthermore, they are also valuable starting points for designing novel drugs against several diseases ranging from cancer to viral infections. Here, we have characterized a small molecule inhibitor of METTL3 by protein crystallography, a biochemical binding assay, and a battery of cellular assays. Our METTL3 inhibitor **UZH1a** shows high-nanomolar potency in the biochemical assay, good selectivity against a panel of protein methyltransferases and kinases, and is active in cells. Importantly, METTL3 chemical inhibition induced apoptosis in AML MOLM-13 cells, while osteosarcoma U2Os cells and the embryonic kidney cell line HEK293T seem to be less reliant on cellular m^6^A level for their survival. These findings highlight that for AML therapy, inhibition of methylation alone may be sufficient to cause cell apoptosis and that the scaffolding function of METTL3 does not necessarily have to be targeted. The crystal structure of METTL3/14 in complex with **UZH1a** has revealed its binding mode and will serve as the basis for the development of more potent inhibitors. Future work will focus on the implications of METTL3 inhibition in various disease models.

## Acknowledgements

The authors are grateful to Beat Blattmann for his help with setting up protein crystallization trials. We thank the staff at the Swiss Light Source (Paul Scherrer Institute) for their support with data collection. We thank the Swiss National Supercomputing Centre (CSCS) in Lugano for providing the computational resources. We also thank Dr. Endre Laczko and Dr. Stefan Schauer from Functional Genomics Center Zurich for their help with UPLC-MS/MS analysis. We thank Dr. Ilia Sergachev for his help with image analysis. We thank Dr. Paweł Śledź, Dr. Katherine Rollins, Dr. Yaozong Li, Prof. Dr. Marianne Hürzeler, and Dr. Claude Schärer for interesting discussions. We also thank Dr. Katherine Rollins for reading the manuscript and very useful suggestions for improving its clarity. This work was supported financially by the Swiss National Science Foundation (E.M.-O., grant CRSK-3_190825; A.C., Excellence grant 310030B-189363) and the Swiss Cancer Research foundation (E.M.-O., A.C., grant KFS-5016-02-2020).

## Materials

HEK293T cells were obtained from ATCC. MOLM-13 cells were a gift from Prof. W. Wei-Lynn Wong (University of Zurich). U2Os cells were a gift from Prof. Yang Shi (Harvard Medical School). Dulbecco’s modified essential medium (DMEM) supplemented with 4.5 g/L glucose and GlutaMAX™, RPMI 1640 medium, Opti-MEM® medium, Dulbecco’s phosphate buffered saline (DPBS), Gibco™ Fetal Bovine Serum, Gibco™ Penicillin-Streptomycin, 0.05% Trypsin-EDTA, nuclease-free water, Lipofectamine® RNAiMAX, FITC-conjugated annexin V, and TO-PRO™3 Ready Flow™ reagent were purchased from Thermo Fisher Scientific (Waltham, MA). Cell culture treated dishes, flasks, and multiwell plates were obtained from Corning Inc. (Corning, NY). CellTiter-Glo® 2.0 cell viability assay kit was purchased from Promega (Madison, WI). Human methylcellulose complete medium (HSC003) was purchased from R&D Systems (Minneapolis, MN). Sera-Mag Oligo(dT)-coated magnetic particles and Hybond-N+ membranes were obtained from GE Healthcare (Chicago, IL). GENEzol™ reagent was purchased from Geneaid Biotech (New Taipei, Taiwan). RNeasy mini kit was obtained from Qiagen (Hilden, Germany). Human METTL3 specific siRNAs and non-targeting siRNA as a negative control were purchased from Microsynth (Balgach, Switzerland). METTL3-siRNA-1 sense: 5′-GCACAUCCUACUCUUGUAAdTdT-3′, METTL3-siRNA-2 sense: 5’-GGAGAUCCUAGAGCUAUUAdTdT-3’, METTL3-siRNA-3 sense: 5’-GACUGCUCUUUCCUUAAUAdTdT-3’, NC-siRNA sense: 5’-AGGUAGUGUAAUCGCCUUGdTdT-3’.^35,36^ NEBuffer™ 2 and nucleoside digestion mix were obtained from NEB (Ipswich, MA). Adenosine and N^6^-methyl adenosine (m^6^A) were purchased from Sigma-Aldrich (St. Louis, MO) and Chemie Brunschwig (Basel, Switzerland), respectively, as LC/MS standards. Anti-METTL3 rabbit and anti-β-actin mouse antibodies were from Abcam (Cambridge, UK), IRDye^®^ 800CW goat anti-rabbit IgG and IRDye^®^ 680RD donkey anti-mouse IgG secondary antibodies were obtained from Li-COR Biosciences (Lincoln, NE). The protease inhibitor cocktail tablets were purchased from Roche (Basel, Switzerland). Propidium iodide was purchased from Chemie Brunschwig (Basel, Switzerland). SurePAGE 12% Bis-Tris polyacrylamide gels were obtained from GenScript (NJ).

## Methods

### Synthetic procedures

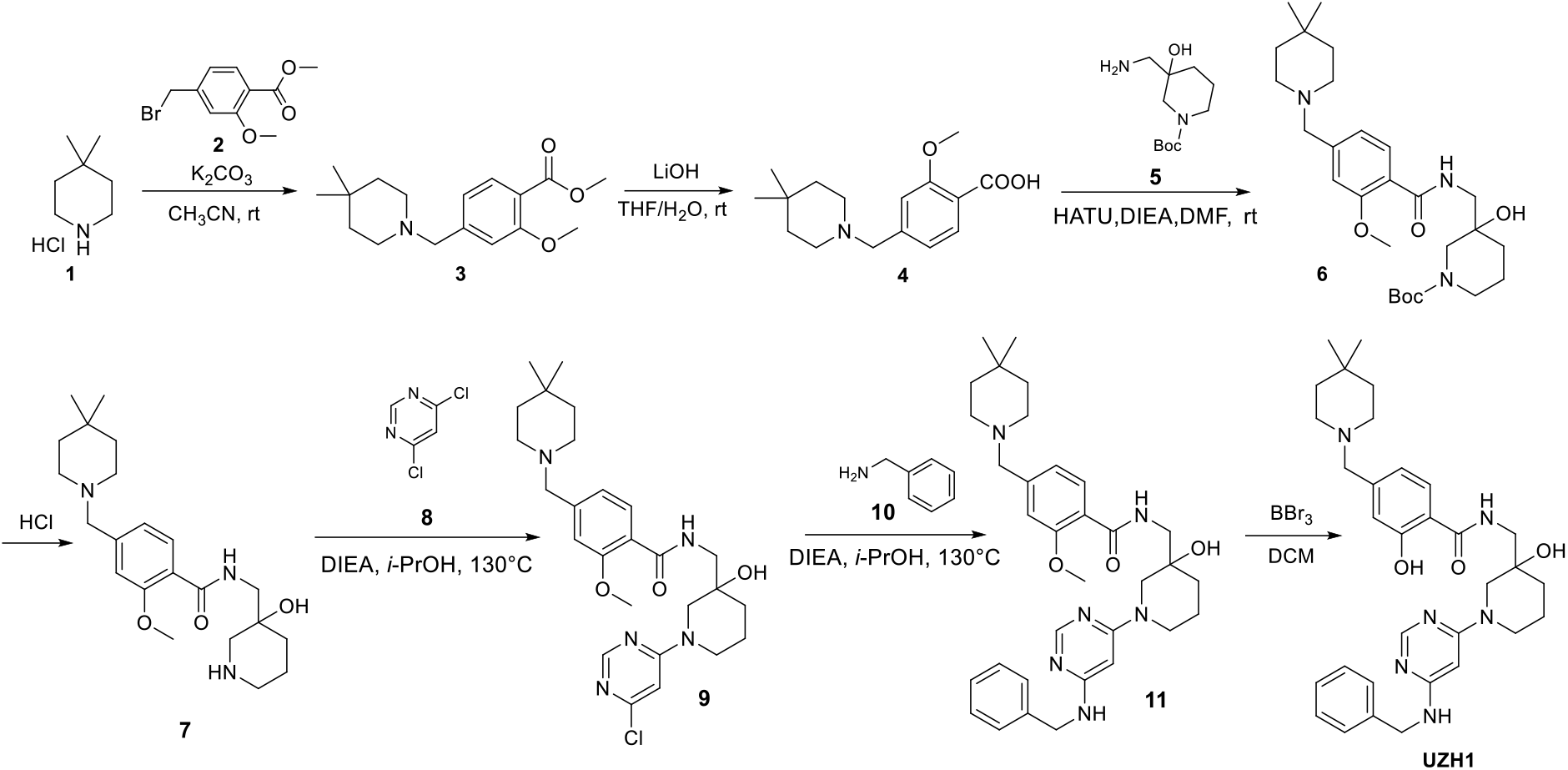

#### Synthesis of compound 3

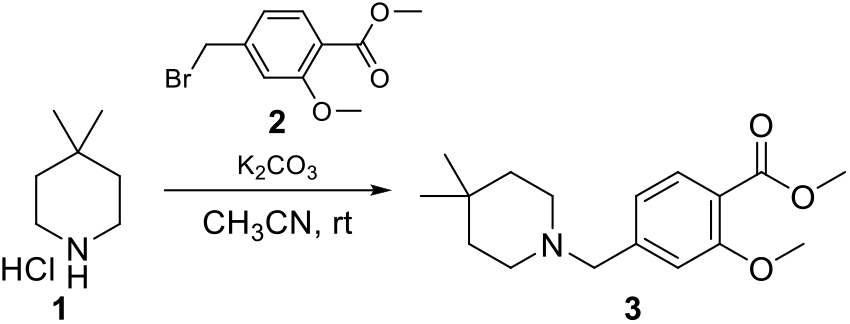

To a solution of 4,4-dimethylpiperidine hydrochloride (350 mg, 2.34 mmol) in CH_3_CN (10 ml) was added methyl 4-(bromomethyl)-2-methoxybenzoate (666 mg; 2.57 mmol) and K_2_CO_3_ (646 mg, 4.68 mmol), then the mixture was stirred at 25 °C for 16 h. The mixture was washed with water (20 mL) and extracted with EtOAc (20 mL × 3), and combined organic layers were dried over anhydrous Na_2_SO_4_, filtered, and concentrated. The residue was purified by column chromatography on silica gel (0-10% EtOAc in petroleum ether) to give methyl 4-((4,4-dimethylpiperidin-1-yl)methyl)-2-methoxybenzoate as a white solid (500 mg, yield: 73%).

#### Synthesis of compound 4

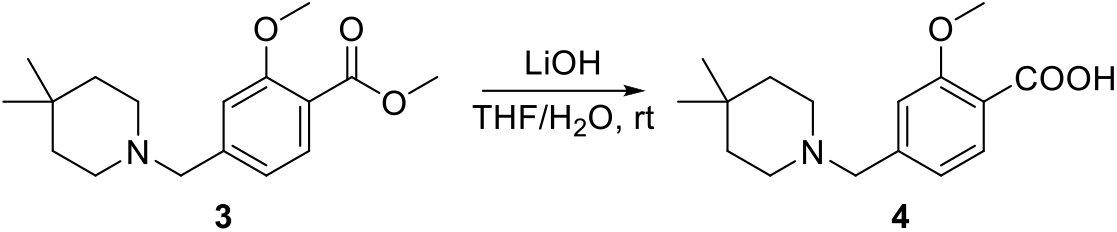

To a solution of methyl-4-((4,4-dimethylpiperidin-1-yl)methyl)-2-methoxybenzoate (500 mg, 1.72 mmol) in THF/H_2_O (10 mL; 5/1) was added LiOH (123 mg, 5.15 mmol), the mixture was stirred at 25 °C for 16 h. The mixture was adjusted by 1 M HCl to pH 3, then extracted with EtOAc (20 mL × 3), and combined organic layers were dried over anhydrous Na_2_SO_4_, filtered and concentrated to give desired product 4-((4,4-dimethylpiperidin-1-yl)methyl)benzoic acid as a white solid (470 mg, crude).

#### Synthesis of compound 6

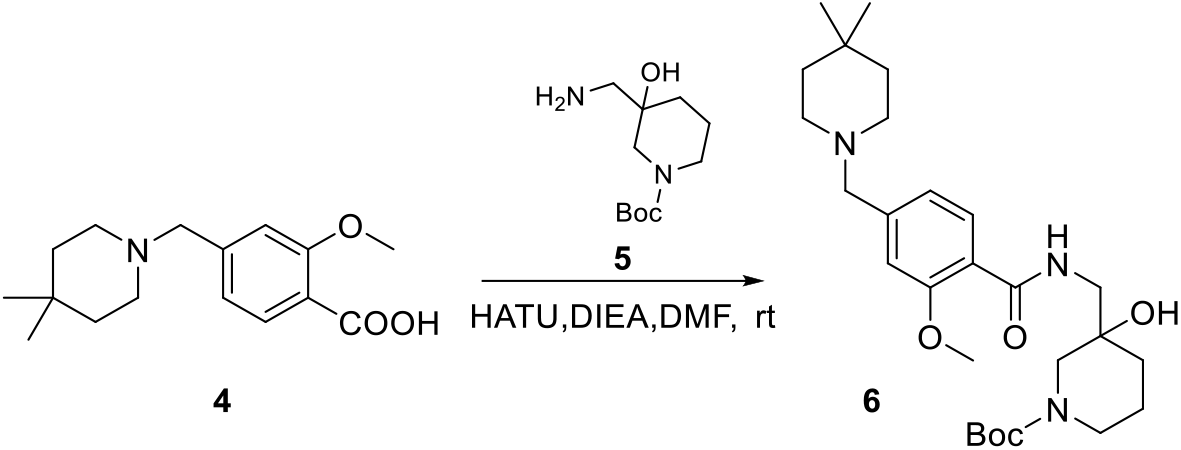

To a solution of 4-((4,4-dimethylpiperidin-1-yl)methyl)-2-methoxybenzoic acid (470 mg, 1.69 mmol) in DMF (10 mL) was added *tert*-butyl 3-(aminomethyl)-3-hydroxypiperidine-1-carboxylate (390 mg, 1.69 mmol) and HATU (773 mg, 2.03 mmol) and DIEA (0.56 mL; 3.39 mmol), then the mixture was stirred at 25 °C for 16 h under N_2_. The mixture was concentrated, washed with water (20 mL), extracted with EtOAc (20 mL × 3), and combined organic layers were dried over anhydrous Na_2_SO_4_, filtered and concentrated. The residue was purified by column chromatography on silica gel (0-5% MeOH in DCM) to give the product *tert*-butyl 3-((4-((4,4-dimethylpiperidin-1-yl)methyl)-2-methoxybenzamido)methyl)-3-hydroxypiperidine-1-carboxylate as yellow oil (560 mg, yield: 67%).

#### Synthesis of compound 7

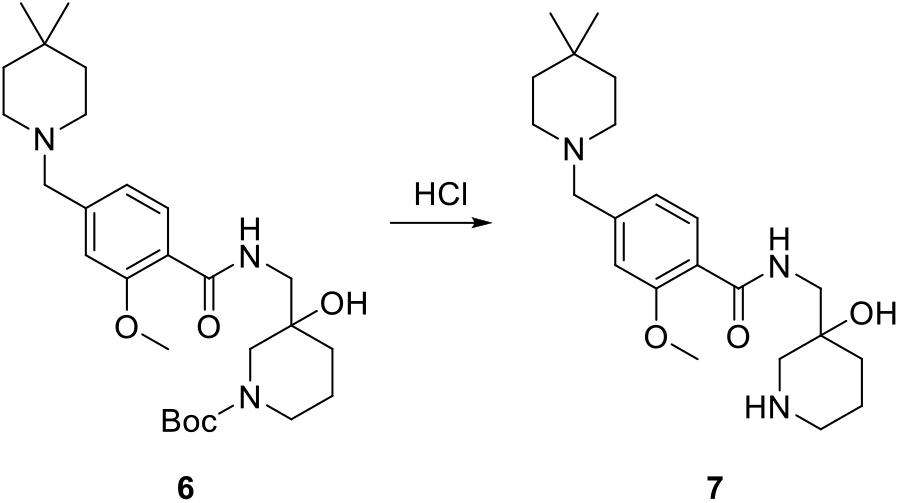

To a solution of *tert*-butyl 3-((4-((4,4-dimethylpiperidin-1-yl)methyl)-2-methoxybenzamido)methyl)-3-hydroxypiperidine-1-carboxylate (560 mg, 1.14 mmol) in DCM (5 mL) was added 3 M HCl in MeOH (1.9 mL; 5.7 mmol), and the mixture was stirred at 25 °C for 16 h. The mixture was concentrated to give desired product 4-((4,4-dimethylpiperidin-1-yl)methyl)-*N*-((3-hydroxypiperidin-3-yl)methyl)-2-methoxybenzamide as yellow oil. (600 mg, crude).

#### Synthesis of compound 9

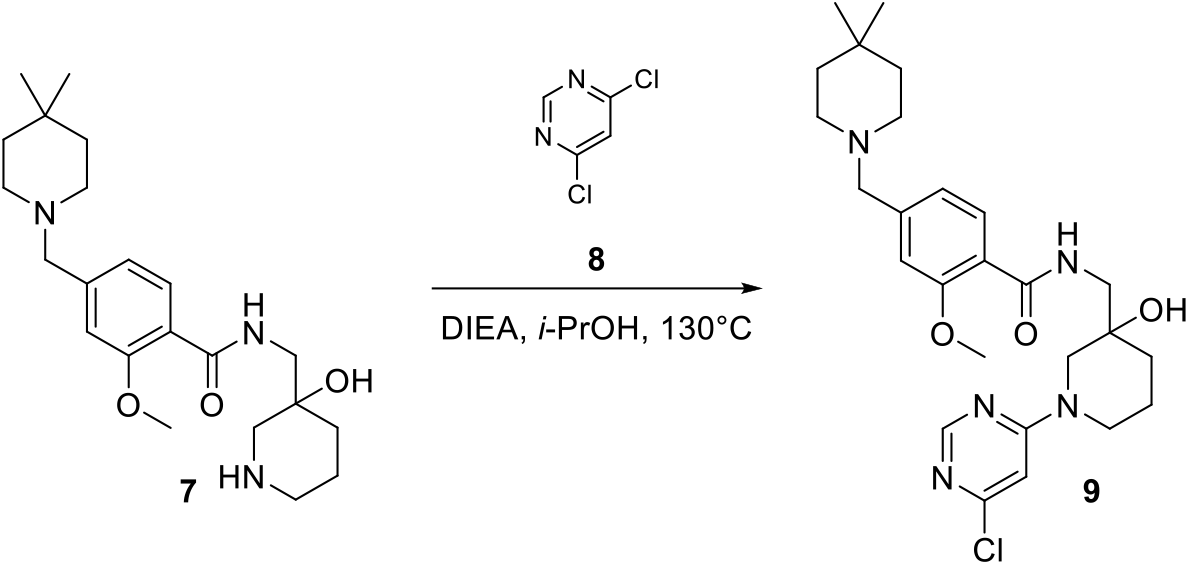

To a solution of 4-((4,4-dimethylpiperidin-1-yl)methyl)-*N*-((3-hydroxypiperidin-3-yl)methyl)-2-methoxybenzamide (680 mg; 1.62 mmol) in *i*-PrOH (10 mL) was added 4,6-dichloropyrimidine (265 mg; 1.78 mmol) and DIEA (0.80 mL; 4.85 mmol). The mixture was stirred at 130°C for 16 h under N_2_. The mixture was concentrated, and the residue was purified by column chromatography on silica gel (0-5% MeOH in DCM) to give the product *N*-((1-(6-chloropyrimidin-4-yl)-3-hydroxypiperidin-3-yl)methyl)-4-((4,4-dimeth ylpiperidin-1-yl)methyl)-2-methoxybenzamide as yellow oil (130 mg, yield: 16%).

#### Synthesis of compound 11

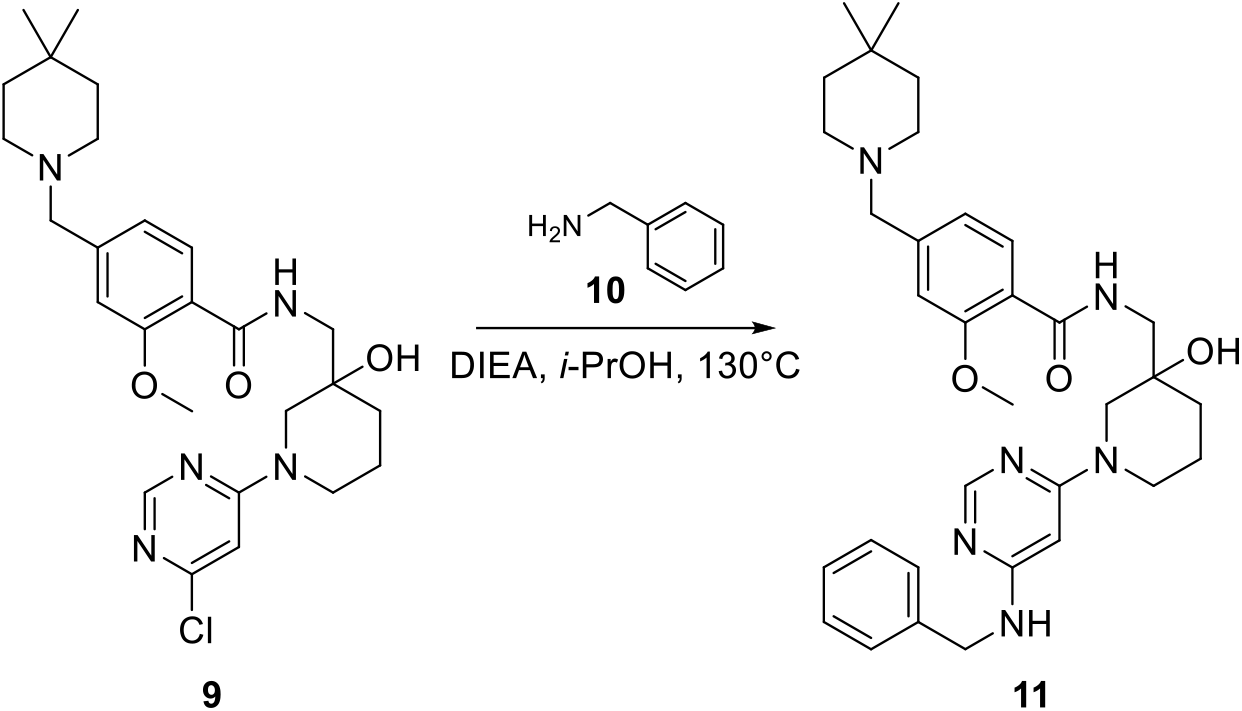

To a solution of *N*-((1-(6-chloropyrimidin-4-yl)-3-hydroxypiperidin-3-yl)methyl)-4-((4,4-dimethylpiperidin-1-yl)methyl)-2-methoxybenzamide (130 mg; 0.26 mmol) in *i*-PrOH (3 mL) was added benzylamine (31 μL; 0.28 mmol) and DIEA (0.12 mL; 0.77 mmol). The mixture was stirred at 130°C for 16 h under N_2_. The mixture was concentrated and the residue was purified by column chromatography on silica gel (0-8% MeOH in DCM) to give the product *N*-((1-(6-(benzylamino)pyrimidin-4-yl)-3-hydroxypiperidin-3-yl)methyl)-4-((4,4-dimethylpiperidin-1-yl)methyl)-2-methoxybenzamide as yellow oil (75 mg, yield: 50%).

#### Synthesis of compound UZH1(racemic)

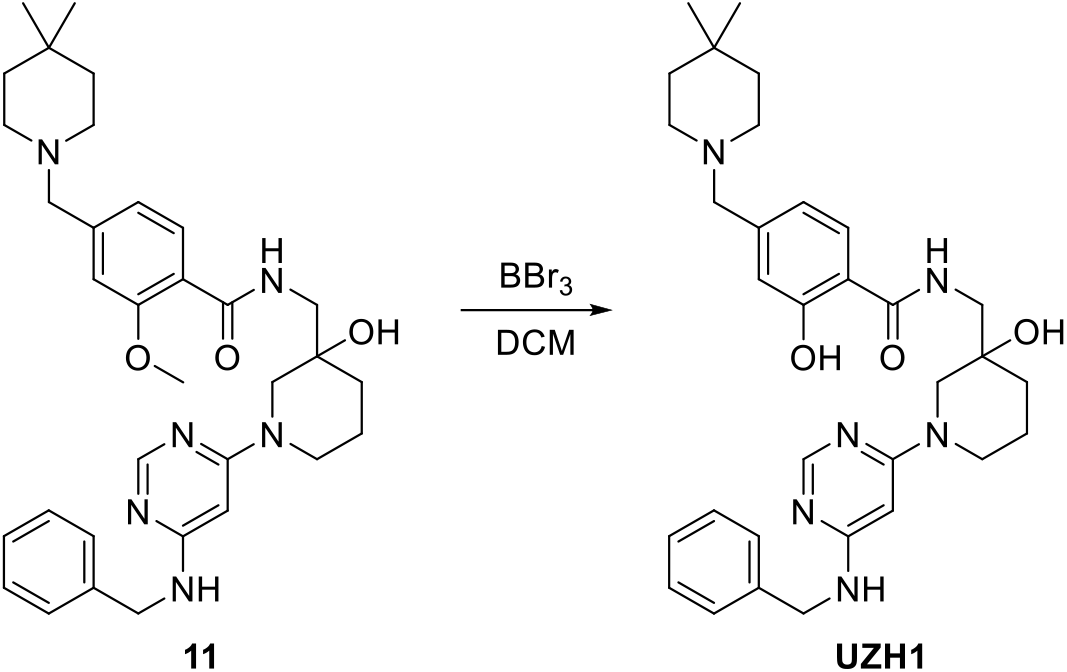

To a solution of *N*-((1-(6-(benzylamino)pyrimidin-4-yl)-3-hydroxypiperidin-3-yl)methyl)-4-((4,4-dimethylpiperidin-1-yl)methyl)-2-methoxybenzamide (75 mg; 0.13 mmol) in DCM (2 mL) was added 1 M BBr_3_ in DCM (0.39 mL; 0.39 mmol) at 0°C. The mixture was stirred at 25°C for 16 h under N_2_. The mixture was concentrated and purified by HPLC to give the desired product *N*-((1-(6-(benzylamino)pyrimidin-4-yl)-3-hydroxypiperidin-3-yl)methyl)-4-((4,4-dimethylpiperidin-1-yl)methyl)-2-hydroxybenzamide as a white solid (24 mg; yield: 32%). The separation of **UZH1** enantiomers was conducted *via* preparative HPLC on a chiral column to give **UZH1a** and **UZH1b**, both as a white solid (purity over 99.5%). It is important to note that the cellular potency of the compounds was crucially affected by their purity, and high degree of purity was required to observe the difference in cellular toxicity between **UZH1a** and **UZH1b**.

### UZH1a

LCMS (ESI): m/z 559 (M+H)^+^

^1^H NMR (400 MHz, CD_3_OD) *δ* 7.95 (s, 1H), 7.82 (d, *J* = 8.0 Hz, 1H), 7.34 – 7.06 (m, 5H), 6.93 – 6.73 (m,2H), 5.58 (s, 1H), 4.37 (s, 2H), 3.61 – 3.36 (m, 8H), 2.50 (s, 4H), 1.85 - 1.72 (m, 2H), 1.71 – 1.57 (m, 2H),1.46 – 1.37 (m, 4H), 0.93 (s, 6H).

### UZH1b

LCMS (ESI): m/z 559 (M+H)^+^

^1^H NMR (400 MHz, CD3OD) *δ* 7.95 (s, 1H), 7.82 (d, *J* = 8.0 Hz, 1H), 7.34 – 7.06 (m, 5H), 6.93 – 6.73 (m,2H), 5.58 (s, 1H), 4.37 (s, 2H), 3.61 – 3.36 (m, 8H), 2.50 (s, 4H), 1.85 - 1.72 (m, 2H), 1.71 – 1.57 (m, 2H),1.46 – 1.37 (m, 4H), 0.93 (s, 6H).

### Protein crystallization

The protein crystals of METTL3_354-580_-METTL14_106-396_ were obtained as previously described.^25^ Briefly, purified METTL3_354-580_-METTL14_106-396_ complex was diluted to 5 mg/mL in 10 mM Tris-Cl pH 8.0, 200 mM NaCl. Crystals were obtained using the hanging drop vapor diffusion method by mixing 1 mL complex solution with 1 mL reservoir solution containing 20% PEG 3350 and 400 mM of magnesium acetate. The soaking experiment was carried out by transferring crystals to a 1 μL drop containing 100 mM compound directly dissolved in the buffer containing 30% PEG 3350, and 400 mM magnesium acetate. After 16 h of incubation at 22°C, the crystals were harvested and flash-frozen in liquid nitrogen.

### Data collection and structure solution

Diffraction data were collected at the PXIII beamline at the Swiss Light Source (SLS) of the Paul Scherrer Institute (PSI, Villigen, Switzerland) and processed using XDS as previously described.^30^ The crystal structures were solved by molecular replacement by employing the 5L6D structure as the search model in the Phaser program^37^ (Phenix package). In the crystals not subjected to soaking, clear electron density for product cofactor S-adenosyl-ʟ-homocysteine (SAH) is visible. Therefore, in this soaking experiment setup test compounds competed with SAH for the *S*-adenosyl-ʟ-methionine (SAM) binding site. In the crystal structure of **UZH1a**, the electron density due to the homocysteine part of SAH was no longer visible. All of the crystallographic models were constructed through iterative cycles of manual model building with COOT^38^ and refinement with phenix.refine.^39^ Default XYZ (reciprocal-space), XYZ (real-space), individual B-factors and occupancies refinement parameters appropriate for the resolution range were utilized. During the first run of the refinement update water was used in phenix.refine followed by addition of the missing water molecules manually.

X-ray data collection and refinement statistics table

| PDB ID | 7ACD |
| --- | --- |
| Compound | UZH1a |
| **Data Collection** |  |
| Space group | P3<sub>2</sub>21 |
| Cell dimension a, b, c (Å) | 63.8, 63.8, 224.92 |
| Cell dimension $\alpha$, $\beta$, $\gamma$ (°) | 90, 90, 120 |
| Resolution (Å) | 44.98 (2.5) |
| Unique reflections* | 19248 (3051) |
| Completeness* | 99.9 (99.8) |
| Redundancy* | 9.72 (9.96) |
| R<sub>merge</sub>* | 19.5 (139.2) |
| CC (1/2) | 99.6 (60.9) |
| I/σI | 11.33 (1.7) |
| **Refinement** |  |
| R<sub>work</sub>/R<sub>free</sub> | 0.1882/0.2259 |
| RMSD bond (Å) | 0.009 |
| RMSD angle (°) | 1.22 |
| B-factor (Å<sup>2</sup>) ** | 45.26/42.22/44.78 |
| Ramachandran Favored | 97.16 |
| Ramachandran Allowed | 2.61 |
| Ramachandran Disallowed | 0.23 |

\*Statistics for the highest resolution shell is shown in parentheses.
\*\* P/L/W indicate protein, ligand/ion and water molecules, respectively.

### Reader-based HTRF assay of METTL3 inhibition *in vitro*

Compound potencies were evaluated by using a previously reported METTL3 inhibition assay.^31^ Briefly, the level of m^6^A in the oligoribonucleotide substrate after the reaction catalyzed by METTL3-METTL14 was quantified by measuring specific binding of modified oligoribonucleotide to the m^6^A reader YTHDC1_345-509_ by homogeneous time-resolved fluorescence (HTRF). Tested compounds that inhibit METTL3 decrease the m^6^A level and, thus, reduce the HTRF signal. Response curves were plotted in GraphPad Prism 8.4 and fitted with nonlinear regression “log(inhibitor) vs. normalized response – variable slope”, from which IC_50_ values were determined. The IC_50_ values are given as an average of at least three independent measurements for each compound.

### Selectivity profiling

For kinase inhibition testing, Merck-Millipore KinaseProfiler™ assay (radiometric, activity testing) was performed by Eurofins (Luxembourg). **UZH1a** was tested in single dose mode, in duplicate, at 10 μM, concentration of ATP was 10 μM. The required volume of the 50× stock of **UZH1a** in DMSO was added to the assay well, before a reaction mix containing an enzyme and substrate was added. The reaction was initiated by the addition of ATP. There was no pre-incubation of the compound with the enzyme/substrate mix prior to ATP addition. For further details of each individual assay, please refer to the company’s website.

For protein methyltransferase inhibition testing, HotSpot Methyltransferase Profiling assay was performed by Reaction Biology Corporation (Malvern, PA). **UZH1a** was tested in single dose mode, in duplicate, at 10 μM. Control compounds, SAH (*S*-(5'-adenosyl)-ʟ-homocysteine) or chaetocin, were tested in 10-dose IC_50_ mode with 3-fold serial dilution starting at 100 or 200 μM. Reactions were carried out at 1 μM SAM.

### Caco-2 cell permeability assay

The **UZH1a** intestinal permeability has been evaluated in Caco-2 cell permeability assay by ChemPartner (Shanghai, China) following standard procedures. Briefly, Caco-2 monolayer cells were cultured in Millicell-24® cell culture plates (Merk Millipore; surface area of the membrane = 0.7 cm^2^, V_A_ = 0.8 mL (A-to-B) or 0.4 mL (B-to-A)). The cell permeability of **UZH1a** along with the reference compounds (*i.e.*, erythromycin, metoprolol, and atenolol) was tested at 10 μM in HBSS buffer containing 0.4% DMSO (v/v) final concentration. The compounds were placed either in apical or basolateral chamber, and their concentrations were evaluated in both compartments after the 90-min incubation at 37°C using the API 4000™ LC/MS/MS system (Applied Biosystems, Waltham, MA). The Caco-2 monolayer’s leakiness after the treatment was evaluated by measuring concentrations of a fluorescent dye Lucifer Yellow with low cell permeability (5 μM initial concentration) using a fluorometer (at Ex/Em of 485/535 nm). Lucifer Yellow P_app_ values were lower than 1·10^-6^ cm/s, and transepithelial electrical resistance (TEER) values were higher than 400 Ω·cm^2^, verifying that the Caco-2 monolayers were intact. Mass recovery of **UZH1a** was 82.4% (A-to-B) and 92.4% (B-to-A), indicating low non-specific adsorption to the assay chambers. Compound permeability was evaluated in duplicates.

Compound permeability was calculated according to the following equation, where VA is the volume in the acceptor well, area is the surface area of the membrane and time is the total transport time in seconds:

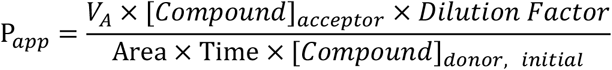

Lucifer Yellow permeability was calculated according to the following equation:

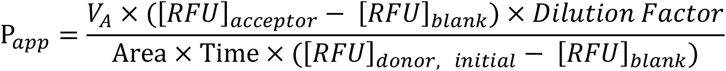

Compound recovery was calculated according to the following equation:

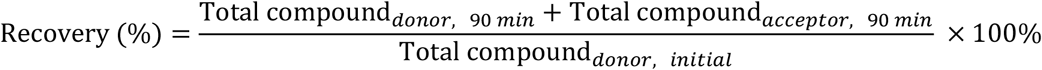

Transepithelial electrical resistance (TEER) was calculated according to the following equation:

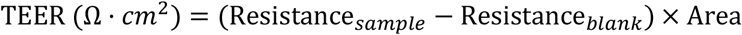

### Cell culture

U2Os and HEK293T cells were maintained in DMEM medium supplemented with 4.5 g/L glucose, GlutaMAX™, 10% Gibco™ FBS, and 1% penicillin/streptomycin (complete medium) in 5% CO_2_ at 37°C in a humidified incubator, with regular passaging twice a week using 1:5 split ratio. MOLM-13 cells were cultured in RPMI 1640 medium containing 10% Gibco™ FBS and 1% penicillin/streptomycin (complete medium) in 5% CO2 at 37°C in a humidified incubator, with maintained cell densities at 0.6 – 2·10^6^ cells/mL. All cell lines have been authenticated by cell line typing (external service provided by Microsynth, Switzerland). All cell lines were tested negative for mycoplasma contamination (PCR-based assay by Microsynth, Switzerland).

### Quantification of m^6^A/A ratio in polyA RNA by UPLC-MS/MS analysis

MOLM-13 cells were seeded into 6 well plates at a density of 1·10^6^ cells/mL in 2 mL of complete RPMI medium. U2Os cells were seeded at 5·10^5^ cells/well in a 6-well plate, whereas HEK293T cells were seeded at 1·10^6^ cells/well. After 24 h, cells were treated with increasing concentrations of **UZH1a** or **UZH1b** in DMSO or DMSO alone as a negative control (final concentration of DMSO 0.5% (v/v)) for indicated time points. Following incubation in cell culture incubator, cells were washed once with PBS, and total RNA was extracted using 0.5 mL of GENEzol™ reagent according to manufacturer's instructions. The final volume of 50 μL of total RNA eluate was subjected to two rounds of purification using 25 μL Sera-Mag magnetic oligo(dt) particles per sample. The polyadenylated RNA was eluted with nuclease-free water in a final volume of 25 μL, and its concentration was determined using NanoDrop. One hundred nanograms of mRNA was digested to nucleosides and dephosphorylated in a one-pot reaction using 0.5 μL of nucleoside digestion mix in 25 μL of total reaction volume for 4 hours at 37°C. The samples were used for UPLC-MS/MS analysis without further purification steps.

Nucleoside UPLC-MS/MS was performed at the Functional Genomics Centre Zurich following previously described procedures with slight modifications.^40,41^ Briefly, the samples were diluted 100-fold in solvent A (0.1% (v/v) formic acid in water), separated using a Waters Aquity M class (Waters) UPLC and detected with a TSQ Quantiva (Thermo Scientific) mass spectrometer by using a nano electrospray ionization (ESI) source in SRM mode. Analytes were separated on an HSS T3 column (particle size 1.8 μm, dimensions 0.15 × 60 mm). The mobile phases used for elution were A 0.1% (v/v) formic acid in water and B 0.1% (v/v) formic acid in acetonitrile at a flow rate of 2 μL/min, column temperature was ambient, sample temperature was 8 °C, injection volume was set to 1 μL, and sample loop volume was 1 μL. The analytes were separated following a gradient of 0 – 4 min 12.5% B, 4 – 4.1 min 99% B, 4.1 – 4.6 min 12.5% B, 6 min 12.5% B. Adenosine and m^6^A eluted at 3.07 min and 3.25 min, respectively. The mass spectrometer was operated using the following parameters: ESI positive ion mode, SRM acquisition mode, capillary voltage 3 kV, source temperature 300 °C, collision energy of 15 and 25 V for adenosine and m^6^A, respectively. Adenosine and m^6^A were detected by monitoring precursor to product transitions of m/z = 268.1 → 136.1 and m/z = 282.1 → 150.1, respectively. Absolute concentrations of adenosine and m^6^A in the samples were determined based on an external calibration curve generated using adenosine (Sigma-Aldrich, 01890-5G) and m^6^A (Chemie Brunschwig, CBLQB-1055-1g) nucleoside standards. The m^6^A/A ratio of compound-treated samples were normalized to the corresponding value of DMSO-treated negative control. All measurements were performed in technical duplicates. Inhibition curves were plotted in GraphPad Prism 8.4 and fitted with nonlinear regression “log(inhibitor) vs. response – variable slope”, from which IC_50_ values were determined.

### RNAi-mediated knockdown of METTL3

For western blotting experiments, HEK293T or U2Os cells were seeded in 6-well plates at a density of 1·10^6^ cells/well in 2 mL of complete DMEM medium, transfected with total 50 nM of METTL3-targeting siRNA mix (at equimolar ratio) or negative control siRNA using Lipofectamine® RNAiMAX according to manufacturer’s instructions, and incubated at 37 °C with 5% CO_2_. The sequences of siRNAs are provided in the materials section. After 72 h, cells were analyzed for METTL3 protein expression using western blotting as described in the western blotting section.

To evaluate the effect of METTL3 knockdown on cell viability, HEK293T or U2Os cells were seeded in 96-well plates at a density of 5·10^3^ cells/well in 100 μL of complete DMEM medium, transfected with 50 nM of METTL3-targeting siRNA mix (at equimolar ratio) or negative control siRNA using Lipofectamine® RNAiMAX according to manufacturer’s instructions, and incubated at 37 °C with 5% CO_2_. After 72 h, cell viability was evaluated as described in cell viability assay section. The assay was carried out with two technical replicates for each treatment and repeated three times on different days.

### Western blotting

For immunodetection of METTL3 expression level after the compound exposure, MOLM-13 cells were seeded at 1·10^6^ cells/mL in a 6-well plate in complete RPMI media. The next day, cells were treated with the indicated doses of compounds. After 16 hours of treatment, cells were collected by centrifugation, washed once with PBS, and resuspended in 200 μL of RIPA buffer (150 mM sodium chloride, 1% (v/v) IGAPAL CA630, 0.5% (w/v) sodium deoxycholate, 0.1% (v/v) sodium dodecyl sulphate, 50 mM Tris-HCl) with added protease inhibitor cocktail (1× final concentration). After incubating for 30 minutes on ice, cell lysates were centrifuged for 15-20 minutes at 16000 × g at 4°C to remove cell debris, and the supernatant containing proteins was collected. The protein concentration was quantified with Pierce™ Coomassie (Bradford) protein assay kit, and 30 μg of protein was loaded per well on a SurePAGE 12% Bis-Tris polyacrylamide gel. Following electrophoresis, the proteins were transferred to methanol-activated low fluorescence PVDF membrane using Trans-Blot® Turbo™ Transfer System (Bio-Rad, Hercules, CA) according to manufacturer's instructions. The membrane was then blocked with 5% non-fat milk and incubated with anti-METTL3 rabbit (1:1000) and anti-β-actin mouse (1:5000) antibodies overnight at 4°C. The next day, the membrane was washed 3 times with PBST (PBS with 0.1% (v/v) Tween-20) and incubated with IRDye^®^ 800CW goat anti-rabbit IgG and IRDye^®^ 680RD donkey anti-mouse IgG secondary antibodies (1:10000) for 2 hours at room temperature. After incubation, the membrane was again washed three times in PBST, and fluorescence signal was detected on Odyssey^®^ CLx Imaging System (LI-COR). The band intensity in each lane was quantified using Image Studio Lite Version 5.2.5 (LI-COR). For the detection of METTL3 protein in U2Os and HEK293T cells following the siRNA-mediated knockdown, the cells were washed with PBS and lysed in 60 μL of lysis buffer. Cell lysates were probed for METTL3 expression as described above.

### Cell viability assay

MOLM-13 cells were seeded in white clear bottom 96-well plates at a density of 2·10^4^ cells/well in 100 μL of complete RPMI medium. HEK293T and U2Os cells were seeded in white clear bottom 96-well plates at a density of 5·10^3^ cells/well in 100 μL of complete DMEM medium. After 24 hours, cells were treated with 11 μL of increasing concentrations of the indicated compounds dissolved in DMSO (final concentration of compounds 2.5 – 160 μM and DMSO 0.5% (v/v)) or DMSO only as a negative control and incubated for 72 h at 37 °C with 5% CO_2_. The assay was carried out with two technical replicates for each concentration and repeated three to four times on different days. Cell viability was determined using CellTiter-Glo^®^ luminescent cell viability assay based on the detection of ATP according to manufacturer’s instructions. Briefly, 100 μL of the CellTiter-Glo^®^ reagent was added to each well and incubated for 10 min at room temperature on an orbital shaker. The bottom of the plate was covered with custom-made white plastic sheet, and luminescence was recorded using a Tecan Infinite 3046 M1000 microplate reader from top. Background luminescence value was obtained from wells containing CellTiter-Glo^®^ reagent and medium without cells. The cell viability was calculated according to the following formula, where *I_compound_* refers to the luminescence intensity of cells treated with different doses of the compounds, *I_DMSO only_* refers to luminescence intensity of DMSO-treated cells, and *I_background_* refers to the luminescence intensity of medium:

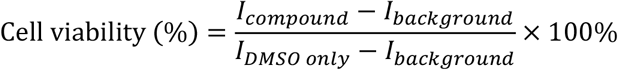

Cell viability curves were plotted in GraphPad Prism 8.4 and fitted with nonlinear regression “log(inhibitor) vs. normalized response – variable slope”, from which IC_50_ values were determined.

### Colony forming unit (CFU) assay

MOLM-13 cells were plated at a density of 1·10^3^ cells/well in 1.5 mL of methylcellulose-containing medium and 200 μL of resuspension medium in 6-well plates in technical duplicates and treated with 5 – 20 μM of **UZH1a** /**UZH1b** in DMSO 0.5% (v/v) or DMSO alone as a negative control. Cells were incubated at 37 °C with 5% CO2 for 10 – 12 days until the untreated cells formed sufficiently large colonies (50 cells or more per colony). Colonies were imaged using EVOS M7000 Imaging System. The experiment was repeated on three different days. Average colony number per treatment was counted manually and the area percentage occupied by cells counted using a scikit-image based python script (https://github.com/el-boo/colony-forming-unit-assay/blob/master/area.py).

### Analysis of cell apoptosis

For flow cytometry analysis of apoptosis, MOLM-13 cells were seeded at 1·10^6^ cells/mL in a 12-well plate and treated with 20 μM of **UZH1a** /**UZH1b** in DMSO 0.5% (v/v) or DMSO alone as a negative control on the same day. After 16 h of treatment, cells were harvested by centrifugation, washed once in PBS and then in 1× homemade annexin binding buffer (HBSS containing Ca^2+^ and Mg^2+^ ions, supplemented with 10 mM HEPES (pH 7.4)). After washing, the cell pellet was resuspended in 1× annexin binding buffer, cells were counted, and their concentration was adjusted to 2·10^6^ cells/mL with 1× annexin binding buffer. One hundred μL of cell suspension (2·10^5^ cells) were stained with 5 μL of FITC-Annexin V and TO-PRO3 (10 μL per sample) on ice for 15 minutes. Cells were diluted with 400 μL of 1× annexin binding buffer, filtered through a cell strainer, and immediately analyzed on a BD FACSCanto™ II Flow Cytometry System (BD Biosciences, San Jose, CA). Cell debris were excluded in SSC-A versus FSC-A plot, and clumps or doublets were gated out in FSC-H versus FSC-A plot. Flow cytometry data was analyzed *via* BD FlowJo software.

### Cell cycle assay

For flow cytometry analysis of cell cycle, MOLM-13 cells were seeded at 1·10^6^ cells/mL in a 12-well plate and treated with 20 μM of **UZH1a** /**UZH1b** in DMSO 0.5% (v/v) or DMSO alone as a negative control on the same day. After 16 h of treatment, cells were harvested by centrifugation, washed once in PBS and then resuspended in 0.5 mL of cold PBS. Cells were fixed for 10 min with 5 mL of ice-cold ethanol added dropwise upon vortexing. After removing the fixation solution *via* centrifugation, the cell pellet was resuspended in 0.5 mL of staining solution (50 μg/mL of propidium iodide, 100 μg/mL of RNAse A, PBS). Cells were incubated on ice for 30 min, filtered through a cell strainer, and immediately analyzed on a BD FACSCanto™ II Flow Cytometry System (BD Biosciences, San Jose, CA). Clumps or doublets were gated out in FSC-H versus FSC-A plot. Flow cytometry data was analyzed *via* BD FlowJo software.

### Statistical analysis

The statistical analysis was performed using GraphPad Prism software. Multiple experimental groups were compared pairwise using the one-way analysis of variance (ANOVA) followed by Tukey’s post-hoc test unless stated otherwise, normality was assessed using Shapiro-Wilk test. The differences between treatment groups were considered statistically significant at *p*-values lower than 0.05.

## Notes

### Competing Interest Statement

The authors have declared no competing interest.

